# Restriction of dietary protein leads to conditioned protein preference and elevated palatability of protein-containing food in rats

**DOI:** 10.1101/210146

**Authors:** Michelle Murphy, Kate Z. Peters, Bethany S. Denton, Kathryn A. Lee, Heramb Chadchankar, James E. McCutcheon

## Abstract

The mechanisms by which intake of dietary protein is regulated are poorly understood despite their potential involvement in determining food choice and appetite. In particular, it is unclear whether protein deficiency results in a specific appetite for protein and whether influences on diet are immediate or develop over time. To determine the effects of protein restriction on consumption, preference, and palatability for protein we assessed patterns of intake for casein (protein) and maltodextrin (carbohydrate) solutions in adult rats. To induce a state of protein restriction, rats were maintained on a low protein diet (5% casein) and compared to control rats on non-restricted diet (20% casein). Under these dietary conditions, relative to control rats, protein-restricted rats exhibited hyperphagia without weight gain. After two weeks, on alternate conditioning days, rats were given access to either isocaloric casein or maltodextrin solutions that were saccharin-sweetened and distinctly flavoured whilst consumption and licking patterns were recorded. This allowed rats to learn about the post-ingestive nutritional consequences of the two different solutions. Subsequently, during a preference test when rats had access to both solutions, we found that protein-restricted rats exhibited a preference for casein over carbohydrate whereas non-restricted rats did not. Analysis of lick microstructure revealed that this preference was associated with an increase in cluster size and number, reflective of an increase in palatability. In conclusion, protein-restriction induced a conditioned preference for protein, relative to carbohydrate, and this was associated with increased palatability.

## Introduction

There is considerable evidence that of the three macronutrients dietary protein is most tightly regulated [1–3]. As such, when presented with diets that differ in macronutrient content, rats will adjust their consumption to ensure that protein intake meets a baseline level [4]. The mechanisms by which these adjustments occur are still not fully understood.

An important outstanding question is whether the drive for protein is immediate and innate or whether there is a role for learning using post-ingestive consequences [5,6]. Some evidence suggests that when protein-restricted a specific appetite for protein arises, similar to the appetite for sodium that arises under conditions of sodium depletion. Rats have been shown to rapidly increase their intake of a number of protein sources when protein-restricted in a manner that precludes using post-ingestive effects to guide their intake [7]. Further research suggested these rapid effects on protein appetite were driven by olfactory cues [8]. However, a large body of evidence indicates that adjustments to protein intake are slow, require experience with each food/diet, and likely involve post-ingestive feedback. For example, when allowed to select between diets that differ in protein content, it takes rats several days to adjust their intake appropriately [9]. This adaptation is more rapid in young rats, although still not immediate, presumably because protein requirements are elevated early in development and positive post-ingestive feedback is enhanced.

The majority of the above studies have assessed food intake and diet selection in home cage tests in which diets are given *ad libitum*. This arrangement does not allow precise monitoring of lick patterns over time. Sophisticated analysis of lick patterns, or lick microstructure, is a key method for assessing palatability of solutions in rodents [10]. As such, when individual licks are grouped into runs based on interlick intervals (termed bursts, clusters and bouts), increases in palatability are associated with longer runs of licking. Importantly, with respect to protein appetite, lick microstructure has not yet been investigated.

Learned shifts in the palatability of protein or protein-containing foods could contribute significantly to increased protein intake under protein-restriction. As a striking example, when rats are sodium-depleted normally aversive concentrations of sodium chloride become highly palatable [11]. Moreover, learning an association between conditioned flavors and intragastric infusions of glucose leads to an increase in palatability of the flavors paired with positive post-ingestive consequences [12,13]. However, increased intake is not always associated with shifts in palatability. For example, rats made deficient in a single essential amino acid increase their intake of the missing amino acid but this is not associated with an increase in palatability [14].

Here, we have used analysis of lick patterns to assess the effect of protein-restriction on intake and palatability of isocaloric protein- and carbohydrate-containing solutions in adult rats. We find that protein-restricted rats, relative to controls, develop a learned preference for protein-containing solutions over carbohydrate and this is associated with an increase in relative palatability.

## Materials and Methods

### Animals

Forty adult male Sprague-Dawley rats were used for experiments (Charles River; >275 g at start of experiment). Twenty-four of these rats were used for the main behavioral experiment and a further sixteen contributed to the food intake data. Rats were group-housed (2-3 per cage) in IVCs with bedding materials as recommended by NC3R guidelines. Temperature was 21 ± 2˚C and humidity was 40-50% with 12h:12h light/dark cycle (lights on at 07:00). Water was available *ad libitum*; chow containing different protein:carbohydrate ratio was available *ad libitum* (details below). All experiments were covered by the Animals [Scientific Procedures] Act (1986) and carried out under the appropriate license authority (Project License: 70/8069).

### Diet manipulations

All rats were initially maintained on standard laboratory chow containing 20% dietary casein. To induce a state of protein restriction in half of the rats, standard chow was switched for experimental diets based on modified AIN-93G that differed in protein:carbohydrate ratio (Table 1). Non-restricted diet (#D11051801, Research Diets, New Brunswick, NJ) contained 20% casein whereas protein-restricted diet (#11092301, Research Diets) contained 5% casein. Body weight data were collected daily throughout the experiments. As rats were group-housed, food intake data were collected by cage and divided by the number of rats in the cage to give an average intake per animal. Conditioning experiments started 2 weeks following diet switch.

**Table 1.** Experimental diets used in study. List of ingredients (upper) and macronutrient breakdown (lower) in control diet (#D11051801; 20% casein) and protein-restricted diet (#D11092301; 5% casein).

|  | **D11051801 (control, 20% casein)** |  | **D11092301 (protein-restricted, 5%)** |  |
| --- | --- | --- | --- | --- |
| **Ingredient** | **g/kg** |  | **g/kg** |  |
| Casein | 200 |  | 50 |  |
| L-Cysteine | 3 |  | 0.75 |  |
| Corn starch | 375.7 |  | 485 |  |
| Maltodextrin 10 | 125 |  | 150 |  |
| Sucrose | 107.1 |  | 107.1 |  |
| Cellulose | 50 |  | 50 |  |
| Soybean oil | 25 |  | 25 |  |
| Lard | 75 |  | 75 |  |
| Mineral mix S10022C | 3.5 |  | 3.5 |  |
| Calcium carbonate | 12.5 |  | 8.7 |  |
| Calcium phosphate, dibasic | 0 |  | 5.3 |  |
| Potassium citrate | 2.48 |  | 2.48 |  |
| Potassium phosphate, monobasic | 6.86 |  | 6.86 |  |
| Sodium chloride | 2.59 |  | 2.59 |  |
| Vitamin mix V10037 | 10 |  | 10 |  |
| Choline Bitartrate | 2.5 |  | 2.5 |  |
| FD&C Yellow dye #5 | 0.05 |  | 0 |  |
| FD&C Red dye #40 | 0 |  | 0.05 |  |
|  | **g (%)** | **kcal (%)** | **g (%)** | **kcal (%)** |
| **Protein** | 18 | 18 | 5 | 4 |
| **Carbohydrate** | 62 | 60 | 76 | 74 |
| **Fat** | 10 | 22 | 10 | 22 |

### Behavioural testing

All testing took place within standard operant chambers (in cm: 30.5 L, 24.1 D, 21.0 H; Med Associates, St. Albans City, VT) equipped with a house light and two bottles. Each bottle was connected to a contact lickometer calibrated to detect individual licks. Licks were recorded on a computer for all sessions as a measure of intake. All sessions lasted for one hour. For one to three days at the start of each experiment, rats were placed in the chambers with 0.2% sodium saccharin in both bottles to familiarize them with the apparatus. Following this, rats underwent a series of conditioning sessions and a preference test. In conditioning sessions, which occurred in a block of 4 days, only one bottle each day was available and was filled with either protein-containing solution (4% casein + 0.21% methionine + 0.2% sodium saccharin + 0.05% flavored Kool-Aid) or an isocaloric carbohydrate-containing solution (4% maltodextrin + 0.2% sodium saccharin + 0.05% flavored Kool-Aid) on alternate days. Methionine was added to the protein-based solution to make up for the relatively low levels of this amino acid that are present in casein [3]. Flavors (cherry vs. grape Kool-Aid) associated with each macronutrient and order of presentation (protein on days 1 and 3 vs. carbohydrate on days 1 and 3) were counter-balanced. In preference test sessions, both bottles and test solutions were available.

### Analysis and statistical methods

Lick timestamp data from all experiments were analyzed in Python. All data files and custom scripts are available as supplemental files and are deposited on Mendeley Data (doi:10.17632/wgd83v3ntb.1). Lick microstructure was analyzed by using interlick intervals to divide licks into clusters [10]. Clusters were defined as runs of licks with no interlick intervals >500 ms.

Body weight data were analyzed using two-way mixed ANOVA with diet as between-subjects factor and day as repeated measure. Food intake data were analyzed with cage as the statistical unit using an unpaired Student’s t-test. Lick data for conditioning days, preference test, and measures of palatability were analyzed using two-way ANOVA with dietary group (non-restricted vs. protein-restricted) as between-subjects factor and solution (casein vs. maltodextrin) as within-subjects factor. On preference test day, protein preference was calculated as licks for casein divided by total licks. Non-restricted vs. protein-restricted rats were compared using unpaired Student’s t-test. For all analyses, α was set at .05 and all tests were two-tailed.

## Results

### Food intake and body weight data across low protein/high protein

First, we assessed whether maintenance on protein-restricted diets affected food intake and body weight of adult rats (Fig. 1). To date, much of the work on protein restriction has used younger rats when protein requirements are greater than in true adulthood. Here, we examined data from rats following the initial dietary manipulation but before conditioning sessions had started so that intake during these sessions did not confound our interpretations.

**Fig. 1.**
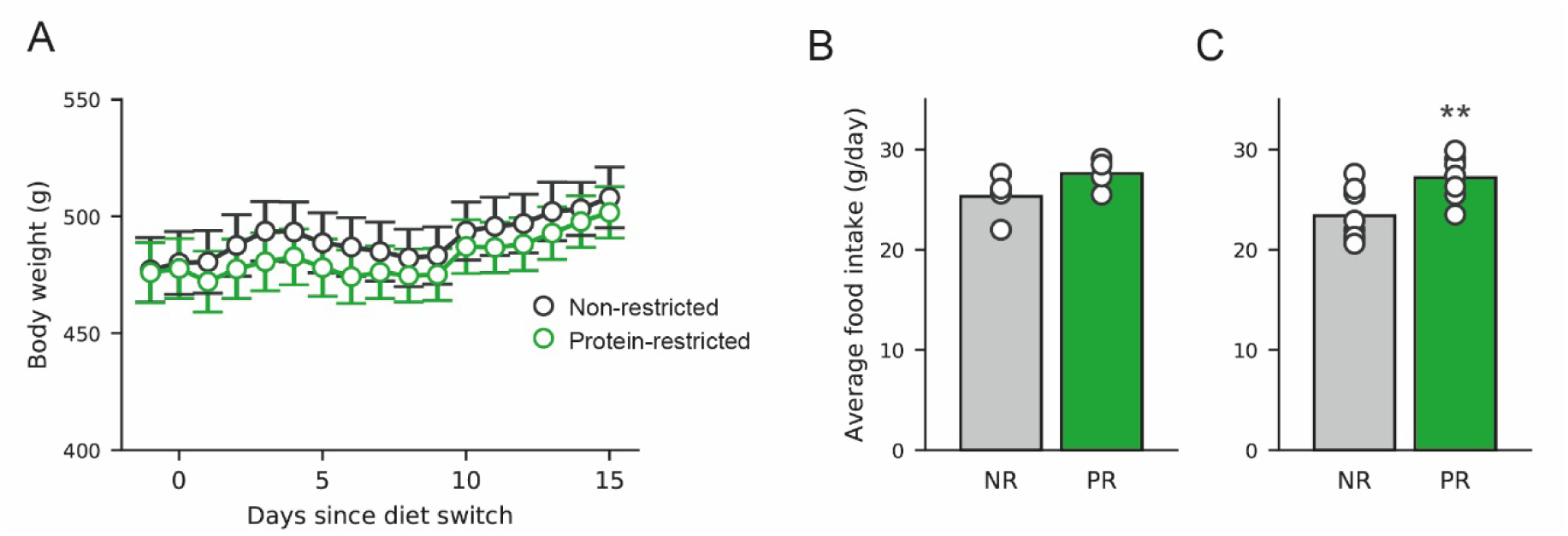
Protein-restricted adult rats increase food intake without changes in body weight. (A) Body weight gradually increases over the course of the experiment in non-restricted (NR; black) and protein-restricted (PR; green) rats with no difference between groups. Data are mean ± SEM. (B) Food intake is greater in protein-restricted rats, relative to non-restricted rats. Intake is shown as grams per day per rat calculated by dividing total daily intake by number of rats in a cage. Bars show mean and data from individual cages are shown as circles. (C) Same data as in (B) supplemented with food intake data from a pilot experiment using rats of comparable age and weight. **, p<0.01 vs. non-restricted rats, [figure = 2 columns]

No difference in body weight was observed between the diet groups over the course of the experiment (Fig. 1A). As such, two-way ANOVA revealed a main effect of Day (F(16,352)=17.371, p<0.001) but no main effect of Diet (F(1,22)=0.115, p=0.738) and no Diet x Day interaction (F(16,352)=0.574, p=0.903).

As rats were group-housed, we obtained food intake data by cage. Food intake data from the eight cages of rats (three rats per cage) that participated in the main study are shown in Fig. 1B and visual inspection suggests a slight increase in intake (hyperphagia) in rats on protein-restricted diet. However, the small number of data points precludes statistical analysis. To address this, we combined this data set with food intake data from a pilot experiment in which an additional eight cages of rats were monitored (two rats per cage) and examined this extended data set (Fig. 1C). Statistical analysis of these data showed that protein-restricted rats did increase their intake of the low protein diet, relative to intake of non-restricted rats (t(15)=3.179, p=0.007). Thus, restriction of dietary protein resulted in hyperphagia without changes in body weight.

### Protein restriction leads to development of preference for protein-containing solutions

Next, we asked whether rats would display a greater preference for protein-containing solutions over carbohydrate-containing solutions when they were protein-restricted. During conditioning sessions, when only one solution or the other was available, we found no significant differences in the amount of consumption between protein-restricted and non-restricted rats although there was a trend for protein-restricted rats to drink more of both solutions than non-restricted rats (Fig. 2). As such, two-way mixed ANOVA revealed a trend towards a main effect of diet (F(1,22)=3.609, p=0.0707) but no main effect of solution (F(1,22)=1.203, p=0.285) and no interaction between diet and solution (F(1,22)=2.087, p=0.163).

**Fig. 2.**
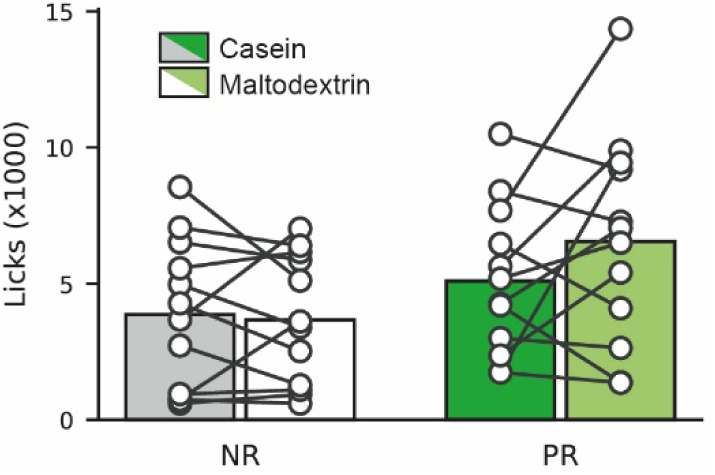
Protein-restricted and non-restricted rats drink similar amounts during conditioning sessions. Rats were given access to distinctly-flavoured casein (4%; 2 × 1 h sessions) or maltodextrin (4%; 2 × 1 h sessions) over four days. Data presented are total licks over both sessions for non-restricted (NR, grey and white bars) and protein-restricted (PR, green bars). Dark bars show casein licks and light/white bars show maltodextrin licks. Bars are mean and circles are data from individual rats. [figure - 1 column]

On day 5, after these four conditioning sessions, rats were given access to both solutions during the same session (Fig. 3). In this session, protein-restricted rats drank more casein than maltodextrin and this elevated intake appeared to occur in the first twenty minutes of the session (Fig. 3A). Furthermore, protein-restricted rats showed a significant preference for casein over maltodextrin whereas non-restricted rats did not (Fig. 3B & 3C). As such, two-way ANOVA revealed that there was a main effect of solution (F(1,22)=7.466, p=0.01216) and an interaction between solution and diet (F(1,22)=11.677, p=0.00247).

**Fig. 3.**
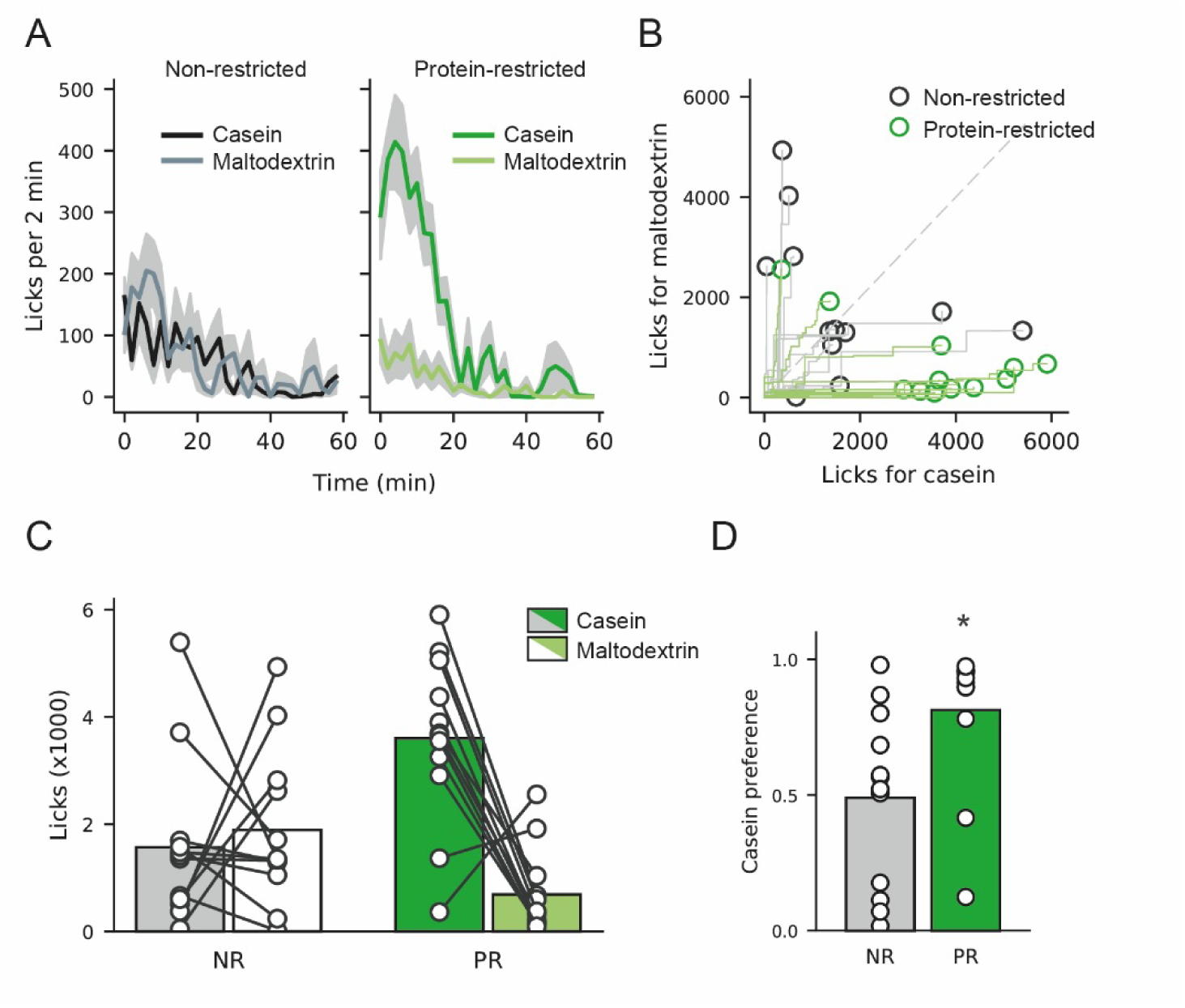
Protein-restricted rats show preference for protein over carbohydrate. After conditioning sessions rats were given access to both casein and maltodextrin solutions within the same session. (A) Time course of licking for casein and maltodextrin during the 1 h preference session. Similar rates of licking are seen for both casein (black line) and maltodextrin (grey line) in non-restricted rats whereas in protein-restricted rats elevated licking is observed to casein (dark green) vs. maltodextrin (light green). This licking predominantly occurs in the first twenty minutes of the session. Lines are mean and shaded area shows SEM. (B) Cumulative licks for casein vs. maltodextrin are shown for individual rats that were non-restricted (grey lines and black circles) or protein-restricted (green lines and circles). Consecutive licks are plotted with casein licks advancing along the x-axis and maltodextrin licks along the y-axis. Dashed grey line at unity represents absence of preference for either solution whereas markers to the right represent casein preference and markers to the left maltodextrin preference. The majority of protein-restricted rats lie to the right of this line indicating protein preference whereas non-restricted rats are evenly distributed. (C) Licks of casein vs. maltodextrin during preference session. Conventions are identical to Fig. 2. (D) Casein preference calculated as casein licks divided by total licks. Protein-restricted rats (green bar) show an increased preference for casein, relative to non-restricted rats (grey bar). Bars are mean and circles are data from individual rats. *, p<0.05 vs. non-restricted rats, [figure = 1.5 columns]

Subsequent analysis of each diet group individually showed that protein-restricted rats licked more for casein than maltodextrin (t(11)=4.630, p=0.0007) but non-restricted rats did not (t(11)=0.458, p=0.656). In addition, we calculated a casein preference score by dividing casein licks by total licks (Fig. 3D) and found that protein-restricted rats showed a greater protein preference, relative to non-restricted rats (t(21)=2.660, p=0.0146).

### Palatability of protein-containing solutions is increased by protein-restriction

Finally, we used analysis of lick microstructure [10] to examine whether the palatability of protein-containing solutions was affected by the state of protein restriction. Lick patterns were divided into clusters, separated by interlick intervals greater than 500 ms. An increased number of licks per cluster is generally thought to reflect increased palatability. We found that the state of protein restriction influenced palatability of casein, relative to maltodextrin (Fig. 4). As such, two-way ANOVA revealed a significant interaction between solution and diet (F(1,22)=7.099, p=0.0142).

**Fig. 4.**
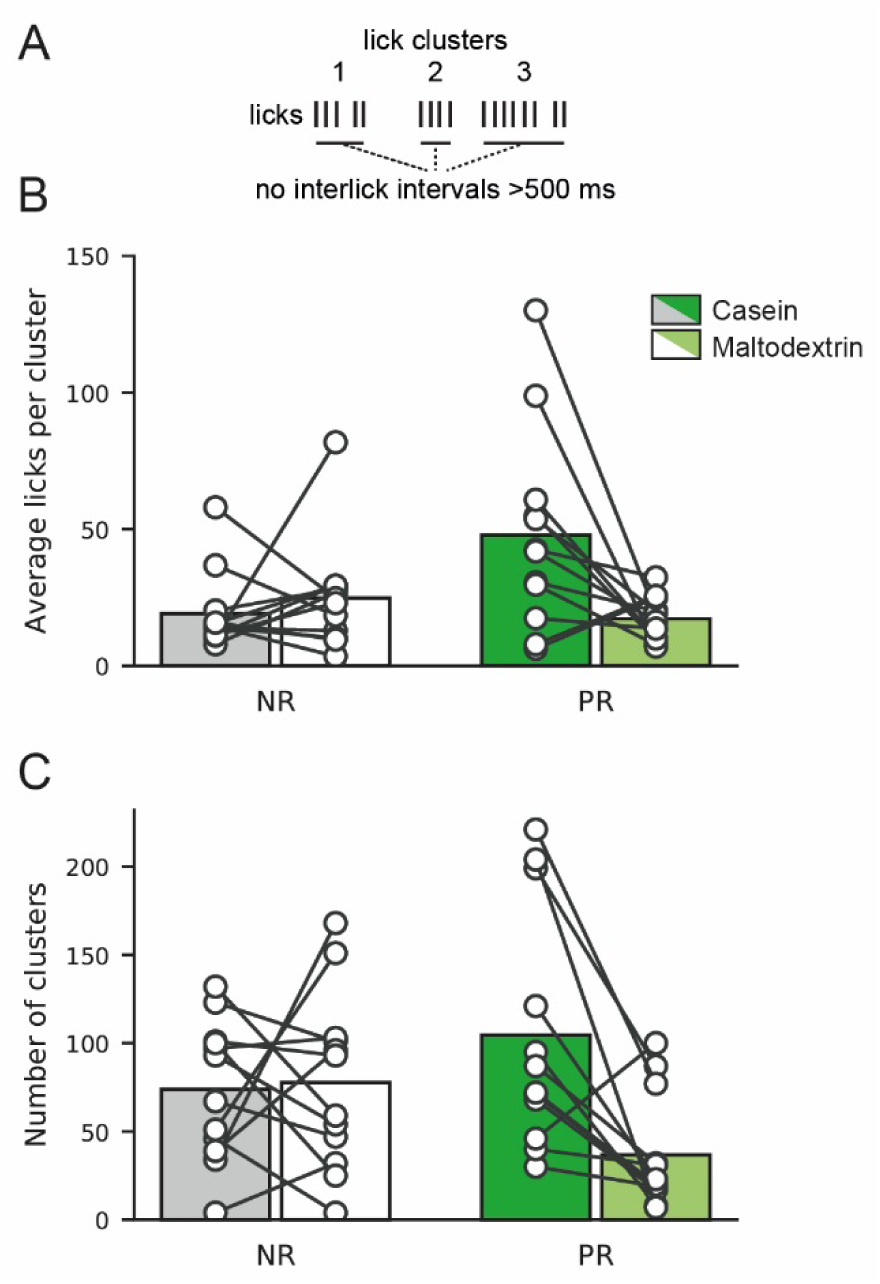
Palatability of protein is enhanced in protein-restricted rats relative to carbohydrate. (A) Schematic showing criteria for defining lick clusters. Licks were grouped into clusters based on having interlick intervals <500 ms. (B) Average licks per clusters in preference test are shown for casein (dark bars) and maltodextrin (light/white bars) for non-restricted (grey/white) and protein-restricted rats (green). Protein-restricted rats show elevated licks per cluster for casein, relative to maltodextrin. Bars are mean and circles are individual rats. (C) Number of clusters is shown for casein and maltodextrin in non-restricted and protein-restricted rats. Conventions are identical to (B). [figure = 1 column]

Further analysis of each diet group separately showed that casein and maltodextrin had similar palatability in non-restricted rats (t(11)=0.761, p=0.4626) but the palatability of casein was elevated relative to maltodextrin in protein-restricted rats (t(11)=2.688, p=0.0211).

In addition, the number of clusters was also influenced by the state of protein-restriction as two way ANOVA revealed a main effect of Solution (F(1,22)=5.677, p=0.0263) and an interaction between Solution and Diet (F(1,22)=7.119, p=0.0140). Analysis of each diet group separately showed that in non-restricted rats there were the same number of clusters for both casein and maltodextrin (t(11)=0.203, p=0.843) whereas protein-restricted rats had an increased number of clusters for casein, relative to maltodextrin (t(11)=3.550, p=0.005).

## Discussion

Here, we examined the effect of protein restriction on development of preference and palatability of protein-vs. carbohydrate-containing solutions. We found that maintenance on a protein-restricted diet resulted in rats developing a preference for protein vs. carbohydrate when given a choice between the two. Moreover, the increase in protein intake was associated with an increase in palatability of the protein-containing solution, relative to the carbohydrate-containing solution.

We monitored food intake and body weight for the two weeks following the change to a protein-restricted diet but before beginning behavioral sessions. Previous studies have found that rats on diets that are moderately low in protein show hyperphagia without weight gain [9,15,16]. In support of these studies, we found that protein-restricted rats increased food intake, relative to controls, without changing their body weight. It is of note, however, that the slight increase in food intake we observe is still far below what is needed to match the protein intake of control, non-restricted rats. In our studies, we used a low protein diet that contained 5% protein whereas other studies using rats have found effects on behavioural and metabolic parameters using diets containing 10% protein [15]. Our choice of 5% was based on pilot experiments, in which we found no effects of 10% protein diet on food intake or conditioned preferences in adult rats (data not shown). The likely explanation for this variation in effective dietary manipulations is different protein requirements during development. Many studies have used late adolescent or young adult rats rather than mature animals and differences in the effects of low protein diets across age and development are well documented [9,17].

In conditioning sessions, over four days rats were given one type of solution - containing either protein or carbohydrate - and lick patterns were monitored. Rats from both dietary conditions drank similar amounts of casein and maltodextrin during these sessions although there was a suggestion (p=0.07) that protein-restricted rats drank slightly more of both solutions than control rats. This may reflect a moderate form of hyperphagia, similar to home cage intake reported above. Interestingly, in the case of these conditioning sessions, when only one solution was available, consumption was increased similarly for the carbohydrate-containing solution meaning that protein-restriction may also generate a hyperphagic response that disregards the macronutrient content of the food on offer.

In the preference test, when rats were given access to both solutions, we found a strong preference towards the protein-containing solution in protein-restricted rats. This preference was not present in control rats. This finding corroborates other work showing that protein-restricted rats can direct their behavior to increase protein intake. In addition, we have extended these previous studies by analyzing the precise temporal patterns of licking to assess how lick macrostructure and microstructure are affected by protein restriction. By analyzing lick microstructure during the preference test, we found that palatability of the protein-containing solutions increased in protein-restricted rats indicating that this might be a mechanism that drives increased intake of protein-containing foods. This situation parallels studies that examined palatability after flavor-nutrient conditioning. When flavored saccharin is paired with intragastric glucose infusions, palatability of the paired flavor is elevated [12]. Our studies used a similar paradigm in which solutions were sweetened with saccharin and distinctly-flavored with Kool-Aid, as is common in studies of flavor-nutrient conditioning [18]. Thus, increased palatability (flavor evaluation) might be a mechanism that drives increased intake by promoting more meals and longer meals.

The presentation of macronutrients in combination with saccharin and flavoring means that we do not know whether the changes in palatability that we observe reflect a change in palatability of individual components of the solution or the combination. When rats are made sodium-deficient, the nutrient itself, sodium, immediately becomes more palatable in an experience-independent manner [11]. This shift is profound as it applies to high concentrations of sodium, which are normally evaluated as aversive in sodium-replete animals. Moreover, sodium-evoked dopamine signals and appetitive behavioral responses to sodium-associated cues also emerge with no experience of sodium in a depleted state [19,20]. Literature suggests that appetite for protein may differ from sodium appetite. For example, when rats are maintained on a diet deficient in a single essential amino acid (lysine), they develop compensatory responses, which increase their intake of lysine, but these responses take ~30 min to emerge and longer if they are required to discriminate between two different amino acids [14]. Interestingly, in this study no evidence of an increase in palatability, assessed by bout size, was observed.

One of the most thought-provoking theories developed to explain the obesity crisis is the protein leveraging hypothesis [21,22]. This theory posits that a steady decrease in the proportion of protein in Western diets occurring over the last few decades has resulted in carbohydrate and fat being overconsumed. The relatively minor role of protein in overall energy intake (generally less than 20%) produces this leveraging ability and means that compensating for even small changes in protein can lead to significant overconsumption of energy from fat and carbohydrate. An important assumption of this hypothesis is that deficiencies in specific nutrients influence our feeding behavior by triggering consumption but that this consumption is indiscriminate and not well-targeted to replenish the nutrient in deficit. Contrary to this assumption, our data suggest that, at least in rats, protein-restriction does recruit mechanisms that enable rats to guide their behavior towards consumption of protein-rich food. However, our studies are far from modelling the human situation and there are numerous important discrepancies to be addressed. First, the level of protein restriction is likely more severe in our protocol than that which most humans in the developed world encounter. Second, the choice of food provided in our studies was limited (protein vs carbohydrate with similar sweetness but distinct flavor) and did not include foods that contained a mixture of macronutrients. Third, the pattern of experience (each solution separately on alternate days) was designed to maximize the ability of rats to discriminate post-ingestive effects and learn about the nutritional value of each solution. In the human situation, where foods contain mixtures of macronutrients and other flavorings, fine discrimination of nutritional consequences of ingestion is likely far more difficult. Moreover, numerous other factors influence our intake such as social setting, cultural norms and access, which may bias us against choosing food stuffs based solely on nutritional outcome. Future studies will attempt to address the ability of rats to develop protein preferences in more challenging situations that better model the human context.

## Acknowledgements

Funding: This work was supported by the Biotechnology and Biological Sciences Research Council [grant # BB/M007391/1]; and the European Commission [grant # GA 631404].

